# *PGSFusion* streamlines polygenic score construction and epidemiological applications in biobank-scale cohorts

**DOI:** 10.1101/2024.08.05.606619

**Authors:** Sheng Yang, Xiangyu Ye, Xiaolong Ji, Zhenghui Li, Min Tian, Peng Huang, Chen Cao

**Affiliations:** Department of Biostatistics, Centre for Global Health, School of Public Health, Nanjing Medical University, Nanjing, Jiangsu 211166, China; Department of Epidemiology, Centre for Global Health, School of Public Health, Nanjing Medical University, Nanjing, Jiangsu 211166, China; Key Laboratory for Bio-Electromagnetic Environment and Advanced Medical Theranostics, School of Biomedical Engineering and Informatics, Nanjing Medical University, Nanjing, Jiangsu 211166, China

**Keywords:** Genome-wide association study (GWAS), polygenic score (PGS), web server, epidemiological application, biobank scale cohort

## Abstract

**Background:** The polygenic score (PGS) is an estimate of an individual’s genetic susceptibility to a specific complex trait and has been instrumental to the development of precision medicine. Clinically, the simplest form of PGS, which is calculated as a weighted sum of variant counts, has been widely applied to conduct disease risk classification. Unfortunately, despite the critical importance of PGS, there are few online resources available to biologists and epidemiologists to calculate PGS in a user-friendly manner.

**Results:** To address this need, we have developed a web server, PGSFusion, that streamlines the construction of PGS using a large variety of methods targeting different epidemiological requirements. PGSFusion included 16 PGS methods in four categories, which are single-trait, annotation-based, multiple-trait, and cross-ancestry. In addition, PGSFusion also utilizes UK Biobank data to provide two kinds of in-depth analyses: i) prediction performance evaluation to display the consistency between PGS and specific traits and the effect size of PGS in different genetic risk groups; ii) joint effect analysis to investigate the interaction between PGS and covariates, as well as the genetic effect size in different subgroups of covariates. PGSFusion automatically identifies the required information in uploaded summary statistics files, provides a selection of suitable methods, and outputs calculated PGSs and their corresponding epidemiological results, all without requiring prior programming knowledge. To demonstrate the function of PGSFusion, we showcase three case studies in different application scenario, highlighting its versatility and values to researchers.

**Conclusions:** Overall, PGSFusion presents an easy-to-use, effective, and extensible platform for PGS construction, promoting the accessibility and utility of PGS for researchers in the field of precision medicine.

## Background

In the past twenty years, genome-wide association studies (GWAS) have identified a large number of significant loci associated with complex traits, including standing height [1, 2], body mass index (BMI) [3], and lipid levels [4], as well as complex diseases, such as coronary artery disease (CAD) [5], type 2 diabetes (T2D) [6], and Alzheimer’s disease (AD) [7]. The proliferation of GWAS has provided a wealth of summary statistics estimated [8] from case-control studies, such as the Psychiatric Genomics Consortium (PCG) [7] and Early Growth Genetics (EGG) Consortium [9], as well as from biobank scale cohorts, such as the UK Biobank (UKBB) [10], China Kadoorie Biobank (CKB) [11], and All of Us [12]. In its simplest form, the polygenic score (PGS) is a weighted sum of the single-nucleotide polymorphisms (SNPs) in an individual’s genotype, where the weights correspond to each SNP’s estimated effect size on a phenotype of interest [13–15]. The widely available summary statistics enable the PGS construction and make it widely used for disease risk stratification, where it is referred to as the polygenic risk score (PRS), with impacts on subsequent clinical applications [16, 17]. However, the efficacy of PGS is hampered by the sizeable differences between GWAS-derived associations from different populations. In particular, this causes PGS constructed from European (EUR) populations to have worse predictive performance when applied to other populations [18]. The introduction of biobank-scale data with multi-ancestry populations is therefore a pivotal step towards enhancing the scalability, transferability, and applicability of PGS.

In the statistical genetics term, based on different assumptions and different estimation strategies, many methods have been proposed as well as compared for their performance among different traits [13, 19–21]. Different traits have different genetic architecture, which results in differing assumptions on the estimated effect size of SNPs. To obtain accurate estimation, emerging PGS methods have been proposed with three different assumptions, including non-model, sparse, and polygenic [13]. For example, Ma *et al.* used DBSLMM to estimate the exposure PGS for 27 exposures in UKBB [22, 23]. Different software uses incompatible summary statistics formats, different reference panel data formats and estimation procedures, and different result formats [24–26]. Besides these hurdles on PGS algorithms, the widespread adoption of PGS by physicians and biologists might face other hurdles, including complex analytic pipelines and the challenge of selecting the most suitable PGS method out of a multitude. It is unfortunate there is a lack of a user-friendly and automatic web server for constructing PGS.

To address this need, we are excited to introduce PGSFusion: a user-friendly webserver that constructs and applies PGS to biobank-scale data (Fig. 1). PGSFusion accepts many different formats of summary statistics and implements a wide range of PGS methods, including single-trait, multiple-trait, annotation-based, and cross-ancestry methods (details in Table 1). Additionally, PGSFusion minimizes the number of parameters that are user-specified, pre-calculates correlation matrices for reference panels from the EUR, African (AFR), and East Asian (EAS) populations of the 1000 Genomes Project (1000GP) [27] and provides annotation information for annotation-based method and validation set in UKBB for parameter selection. Finally, PGSFusion provides further epidemiological analyses for evaluating performance and identifying joint analysis, with all computed PGS and associated analysis figures available to download.

**Fig. 1.**
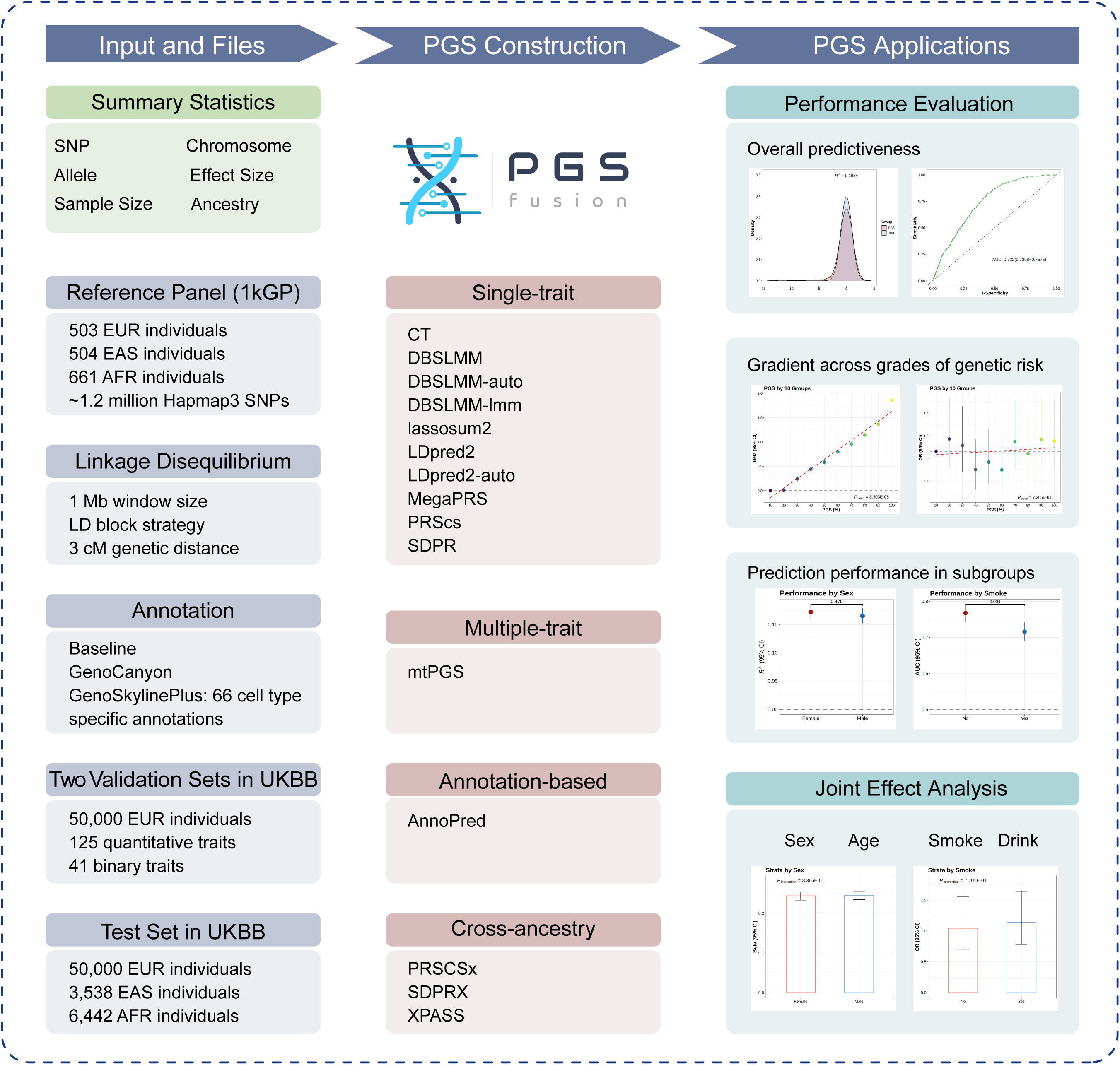
The design of PGSFusion. PGSFusion is a web-based application for constructing and visualizing PGS with an easy-to-use interface. PGSFusion includes 16 PGS construction methods classified into four categories: single-trait, multiple-trait, annotation-based, and cross-ancestry methods. Reference panels containing 503 EUR, 661 AFR, and 504 EAS individuals are drawn from the 1000 Genomes Project. To streamline usability, PGSFusion automatically infers required information from GWAS summary statistics files, provides validation sets from UKBB (EUR population) for 166 common traits (125 quantitative and 41 binary) to automatically select optimal hyper-parameters for six methods, and performs comprehensive epidemiological applications of prediction performance evaluation and joint effect analysis. To construct PGS, the user simply uploads a summary statistics file and selects the available PGS methods, before reviewing visualizations and downloading results via FTP links. Note that methods requiring a validation set should not be applied to summary statistics, including UKBB EUR individuals.

**Table 1.**
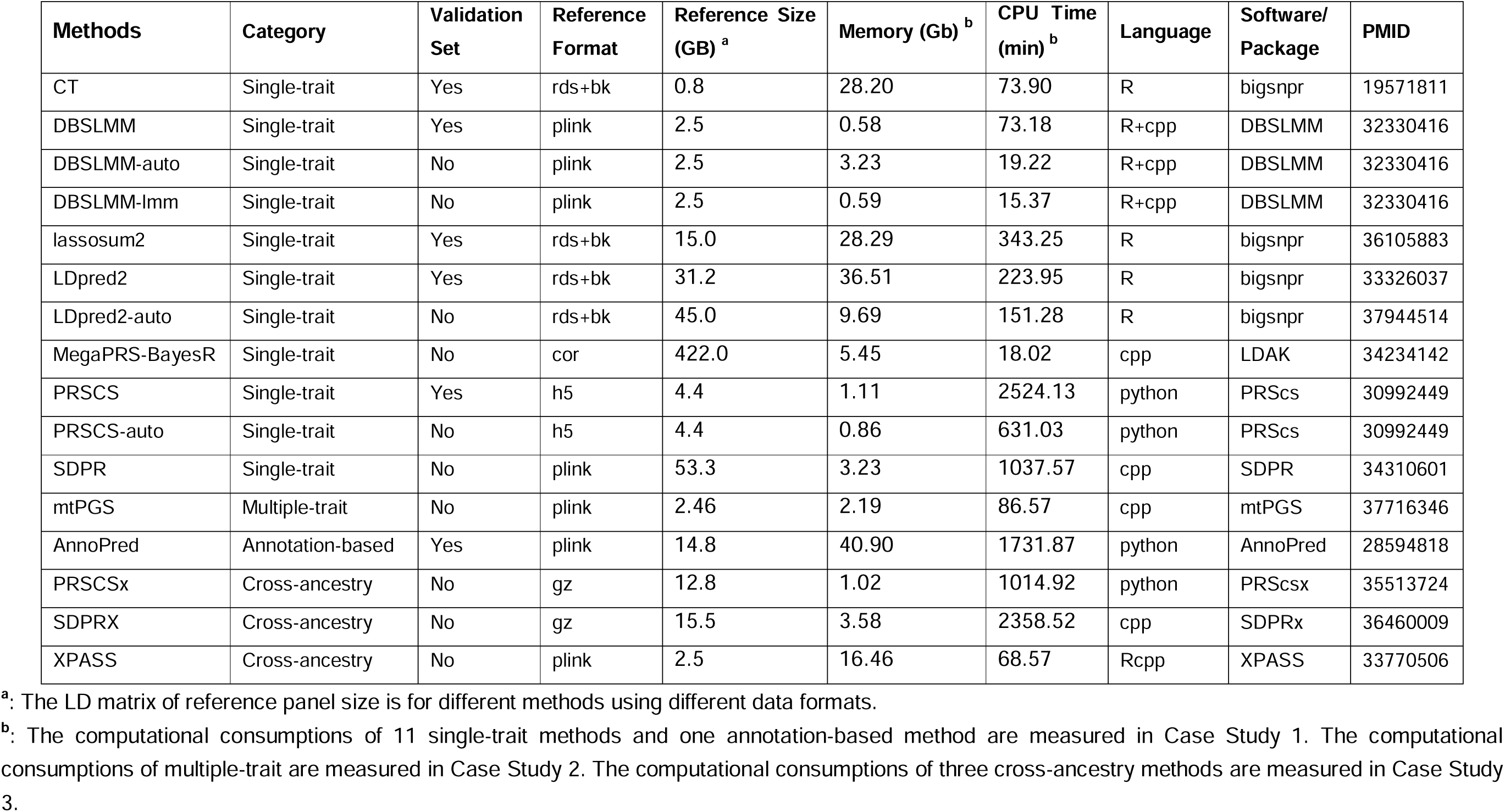
Summary for PGS Construction Methods.

## Results

### Overview

PGSFusion (http://www.pgsfusion.net/) is a cutting-edge webserver that utilizes state-of-the-art web technologies and integrates 16 PGS methods to provide comprehensive and user-friendly analyses for PGS construction (Fig. 1). Users can upload their own summary statistics in a variety of formats, including the GEMMA format, from which PGSFusion automatically identifies the column names of the data required for constructing PGS [28]. To help navigate the different PGS methods and their use cases, we separate the PGS methods included in our web server into four categories: single-trait, multiple-trait, annotation-based, and cross-ancestry (Table 1). To improve accessibility for non-expert users, PGSFusion eliminates manual parameter settings, displays available methods based on summary statistics, and suppresses output information. For methods that select optimal hyper-parameters using a validation set, PGSFusion uses 50,000 EUR UKBB individuals as the validation set (validation set I) and applies these methods to summary statistics from non-UKBB individuals only. When applied to non-UKBB summary statistics, PGSFusion uses an additional 50,000 UKBB individuals as a test set to conduct two further epidemiological applications: i) performance evaluation; ii) joint analysis. PGSFusion provides FTP links for users to download the computed PGS and supports users looking to directly download the PGS application. Specifically, PGSfusion uses 50,000 UKBB individuals as another validation set (validation set II) to determine the coefficients assigned to the constructed PGS and covariates (i.e., sex, age, and top PCs) for the test set.

### Case study 1: PGS construction and applications for AD in PCG

This case study shows how PGSFusion can be used to construct PGS with multiple single-trait methods, while also investigating the prediction performance and joint analysis of the method with the best performance in the EUR test set. After performing QC for SNPs, we obtained 1,223,440 HapMap3 (HM3) SNPs for AD summary statistics. Using LDSC, we estimated the observed heritability of AD to be 0.059 (*P*-value = 7.755×10^-8^), with a genomic inflation factor (*λ*) = 1.099. The Manhattan and Q-Q plots are shown in Additional file 1: Fig. S1.

Given the heritable nature of AD, we used the summary statistics to fit 11 single-trait methods and one annotation-based method to evaluate their prediction performance, computational time, and memory usage (Fig. 2A and 2B). We used the validation set I to tune the default hyperparameters for six methods and utilized the EUR test set to investigate two epidemiological applications of PGS.

**Fig. 2.**
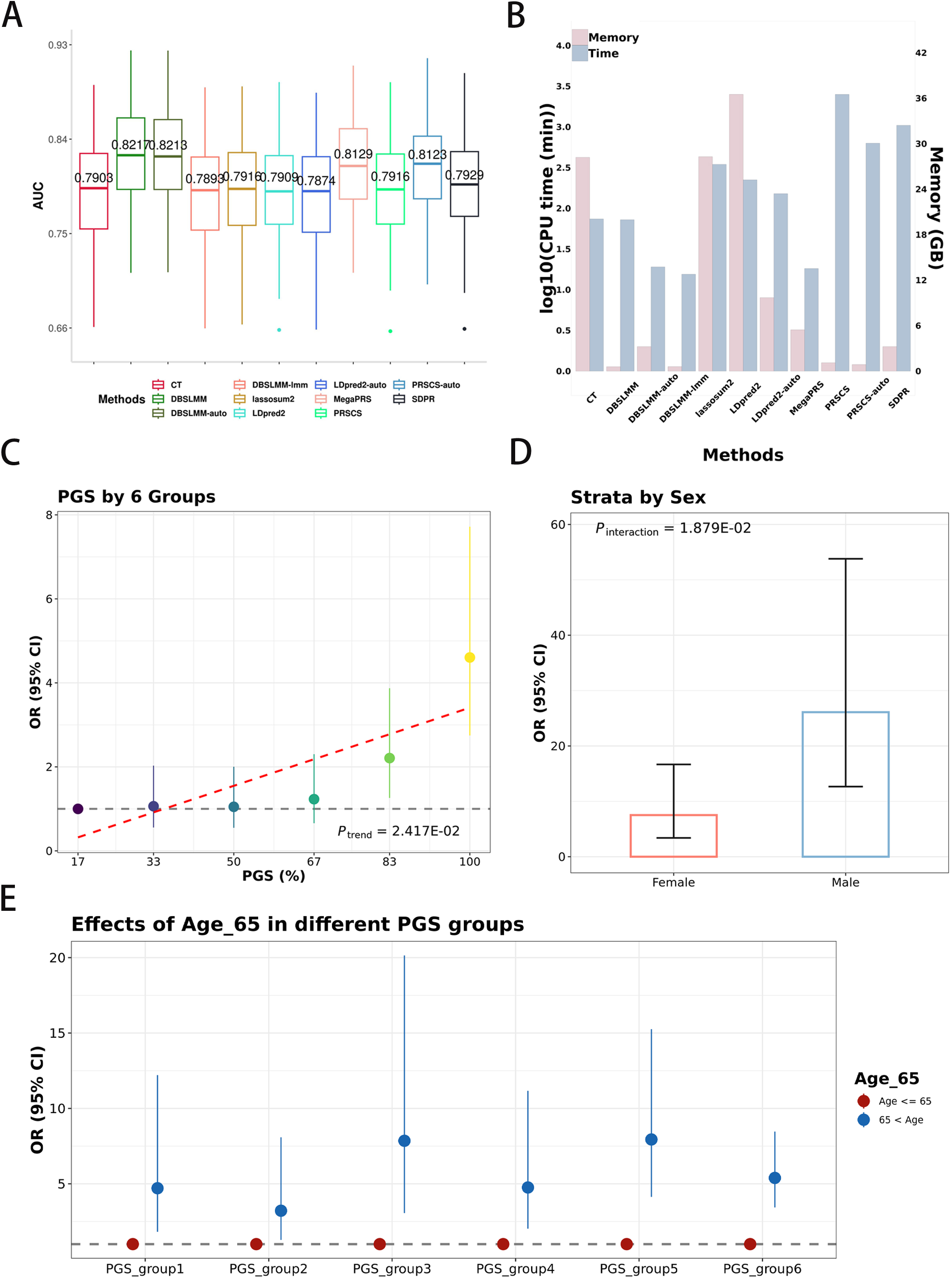
Summary of AD PGS construction and application. A) Box plots showing prediction performance in terms of Pearson correlation coefficient for 11 single-trait methods, estimated via bootstrapping on the EUR test set. B) Computational consumption of four cross-ancestry methods when evaluated on Intel Xeon Platinum 8255C CPU-2.50GHz processors. The two *y*-axes show CPU usage time in log scale (light steel blue) and memory (thistle). C) Forest plot displaying the effect size in different PGS risk groups. Individuals in the EUR test sets were divided into six equal groups according to their risk quantile, and the odds ratios (ORs) between each group and the lowest risk group were computed. Error bars indicate 95% CIs. D) Bar plot showing ORs and 95% CIs for male and female PGS and *P*-values for the effect size of interaction with sex. E) Forest plot displaying the PGS effect size for six age groups.

First, we found that DBSLMM, DBSLMM-lmm, MegaPRS, and PRSCS-auto had comparable prediction performance, with an AUC of approximately 0.822 (Fig. 2A). The AUCs for the other methods were above 0.790. Among the 11 methods, DBSLMM-lmm and MegaPRS were the fastest, with computation times of 15.37 and 18.02 minutes, respectively (Fig. 2B). The most memory-efficient models included the three different DBSLMM models and the two PRSCS models, using approximately 0.60 and 0.86 GB of memory, respectively (Fig. 2B). In contrast, LDpred2, lassosum2, and CT consumed significantly more memory, nearly 30 GB. Specifically, LDpred2, CT, and lassosum2 took 223.45, 73.90, and 343.25 minutes and used 36.51, 28.20, and 28.29 GB of memory, respectively. Although PRSCS and PRSCS-auto were memory-efficient, their MCMC estimation took the longest time, 2524.13 and 631.03 minutes. Considering both computational consumption and prediction performance, we selected the PGS results from DBSLMM or DBSLMM-auto as the primary outcomes and provided the ROC curves for the remaining nine methods in Additional file1: Fig. S2.

Second, dividing the test set into six groups based on PGS, we observed that the effect size significantly increased with higher PGS (*P_trend_* = 2.42×10^-2^) (Fig. 2C and Additional file 2: Table S1). Specifically, OR in the 6^th^ PGS group was 4.605 (95% CI: 2.747 ∼ 7.718, *P*-value = 6.80×10^-9^). The results of other PGS group numbers are shown in Additional file 1: Fig. S3. We also identified a significant interaction between PGS and sex (*P*-value = 1.88×10^-2^) (Fig. 2D). Additionally, the effect size of age was significant across different PGS groups (Fig. 2E and Additional file 2: Table S2). Subgroup analyses for four covariates are provided in Additional file 1: Fig. S4.

### Case study 2: cross-ancestry PGS construction and applications for standing height in BBJ and GIANT

This case study shows how PGSFusion can be used to construct PGS with multiple cross-ancestry methods and conventional single-trait methods in order to compare their performances in cross-ancestry prediction. After applying QC to filter SNPs related to height summary statistics from BBJ and GIANT, we obtain 955,433 and 1,172,056 HM3 SNPs, respectively. Based on LDSC, we estimated the heritability of height in EAS and EUR populations as 0.307 (*P*-value = 0.00) and 0.328 (*P*-value = 0.00), in line with the cross-ancestry genetic correlation of 0.887 computed by PropCorn. The Manhattan plot and QQ-plot are shown in Additional file 1: Fig. S5.

Considering that height is heritable, we used the summary statistics to fit three cross-ancestry and three single-trait methods to compare their prediction performance, computational time, and memory usage (Fig. 3A and 3B). We used the validation set I to tune the default hyperparameter settings for the single-trait methods. We used EAS individuals in UKBB as the test set to evaluate the cross-ancestry prediction performance of each method and to also demonstrate the two epidemiological applications of PGS.

**Fig. 3.**
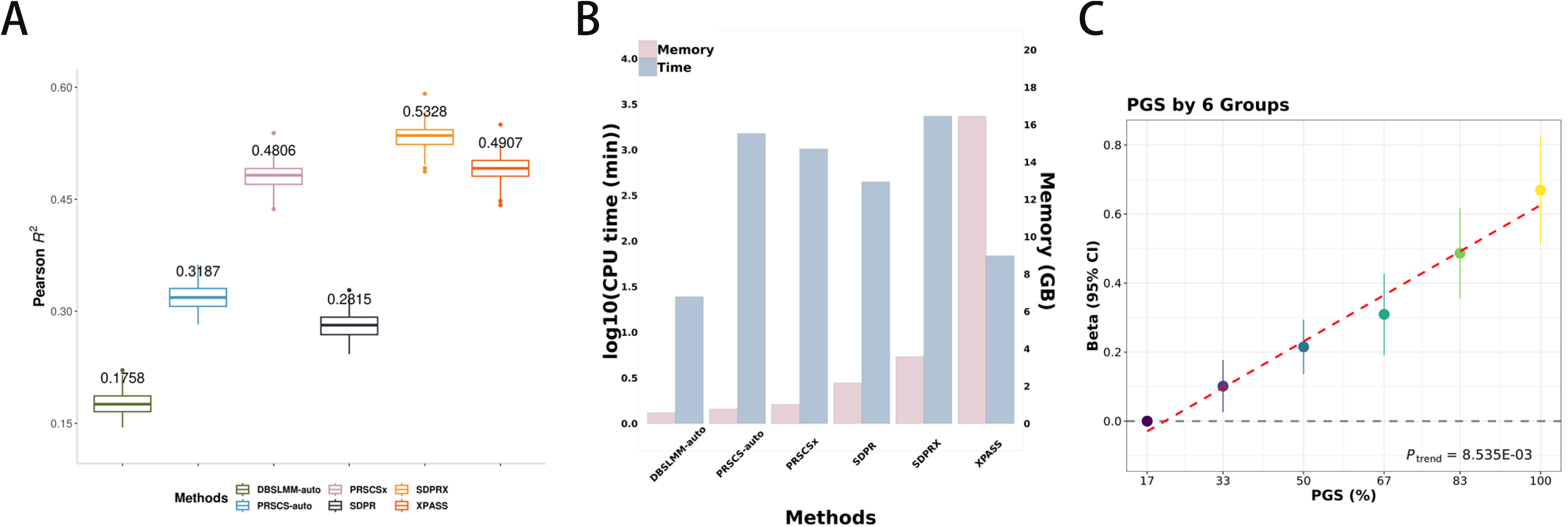
Summary of cross-ancestry standing height PGS construction and application in EAS in the UKBB cohort. A) Box plots showing prediction performance in terms of Pearson correlation coefficients for DBSLMM-auto, PRSCS-auto, PRSCSX, SDPR, SDPRX, and XPASS, estimated via bootstrapping on the EUR test set. B) Computational consumption of the four methods on Intel Xeon Platinum 8255C CPU-2.50GHz processors, including CPU usage time in log scale (light steel blue) and memory (thistle). C) Forest plot of effect size in different score groups. Individuals in the EUR test set were divided into six equal groups according to their PGS quantile, and the betas between each group and the lowest group were computed. Error bars are 95% CIs.

We compare the results of each PGS method as follows. First, the average improvement in prediction performance of cross-ancestry approaches over single-trait methods was 93.83% (mean of *R*^2^ in single-trait methods = 0.259 and mean of *R*^2^ in cross-ancestry methods = 0.501). For methods with similar model assumptions, we compared their cross-ancestry prediction performance pairwise (Fig. 3A and Additional file 1: Fig. S6). For example, when estimating effect sizes using coupled continuous shrinkage priors, PRSCS and PRSCSx give *R*^2^ values of 0.319 and 0.481, respectively, on the EAS test set (averaged over bootstrapping). As another example, using a Dirichlet prior for effect sizes, *R*^2^ values for SDPR and SDPRX were on average 0.282 and 0.533, respectively, on the EAS test set. Similarly, *R*^2^ values for DBSLMM-LMM and XPASS, which assume normally distributed effect sizes, were 0.176 and 0.491 respectively. Second, PRSCSx, XPASS, and SDPRX required 1014.92, 68.57, and 2358.52 minutes to run and used 1.02, 16.46, and 3.58 Gb of memory, respectively (Fig. 3B). Note that the two Bayesian methods (PRSCSx and SDPRX) were, respectively, 14.80 and 34.40 times slower than XPASS.

Next, we considered the PGS with the highest prediction performance, which is constructed using SDPRX. By dividing the test set into six groups using this PGS, we found that the effect size increased across PGS groups (*P_trend_*= 8.54×10^-3^) (Fig. 2C and Additional file 2: Table S3). Note that the effect size in the 6^th^ group was 0.669 (95% CI: 0.511 ∼ 0.827, *P*-value = 2.00×10^-16^). The results for the other test set grouping are shown in Additional file 1: Fig. S7.

### Case study 3: PGS construction and applications for weight in males leveraging HDL

This case study shows how multiple traits can be used to enhance PGS construction, while also showcasing the sex-specific analysis function of PGSFusion. After applying QC to summary statistics, we obtained 1,144,828 and 1,169,070 HM3 SNPs for weight and HDL, respectively. The Manhattan and Q-Q plots are shown in Additional file 1: Fig. S8 with ..1 = 0.999 and 1.710. Using LDSC, the heritability of the two traits was 0.172 (*P*-value = 0.000) and 0.225 (*P*-value = 0.000), with a genetic correlation of −0.328 (*P*-value = 3.04×10^-25^). Given that there was a significant genetic correlation between the traits, we used the mtPGS model to leverage the HDL trait to construct a PGS targeting weight. Using similar assumptions on the effect size distribution, we also compared the prediction performance and computational cost of mtPGS with DBSLMM.

Following [15], we evaluated performance over 100 bootstrap samples with 5,000 EUR individuals per sample. Since DBSLMM did not leverage the HDL trait, mtPGS had the highest prediction performance as expected, with *R*^2^ = 0.022 (Fig. 4A, 4C, and Additional file 1: Fig. S9). The computation time and memory consumed by mtPGS were 86 minutes and 2.19 Gb, respectively (Fig. 4B). Although mtPGS took 4.778 times longer and used 3.778 times more memory than DBSLMM-auto, the prediction performance was improved by 44%. When dividing the test set into five groups by PGS, we found that the effect size was significantly increasing using mtPGS (*P_trend_*= 2.75×10^-2^) (Fig. 4D and Additional file 2: Table S4). We also estimated effect sizes for four variables in each of the five PGS subgroups. Notably, the effect size of smoking formed a U-shaped relationship across the groups (Fig. 4E). While the effect size of smoking in the 3^rd^ group was 0.097 (95% CI: 0.042 ∼ 0.151), those of the 1^st^ and 5^th^ groups were 0.103 (95% CI: 0.052 ∼ 0.153) and 0.145 (95% CI: 0.087 ∼ 0.203), respectively (Additional file 2: Table S5). Results for all PGS subgroups are shown in Additional file 1: Fig. S10, while the subgroup analysis for the remaining two covariates is in Additional file 1: Fig. S11.

**Fig. 4.**
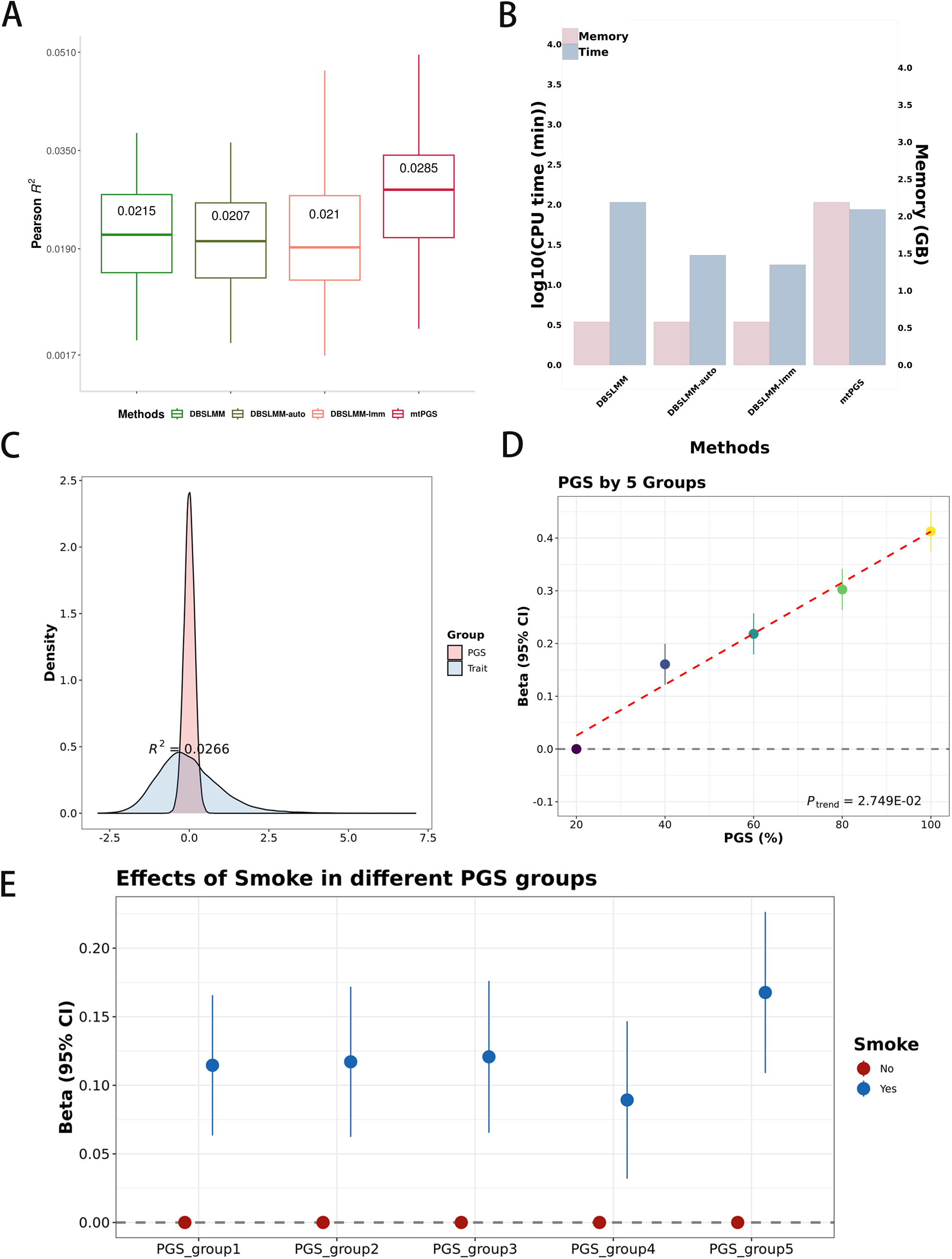
Summary of results from constructing weight PGS while leveraging HDL in males. A) Box plots showing prediction performance in terms of Pearson correlation coefficients for DBSLMM, DBSLMM-auto, DBSLMM-lmm, and mtPGS, estimated via bootstrapping on the EUR test set. B) Computational consumption of the four methods on Intel Xeon Platinum 8255C CPU-2.50GHz processors, including CPU usage time in log scale (light steel blue) and memory (thistle). C) Distribution of trait versus PGS for weight in the EUR test set. D) Forest plot of effect size in different score groups. Individuals in the EUR test set were divided into five equal groups according to their PGS quantile, and the betas between each group and the lowest group were computed. Error bars are 95% CIs. E) Forest plot displaying the effect of smoking on PGS for the five groups.

## Discussion

We have designed and developed PGSFusion to streamline PGS construction and epidemiological analysis, making it more accessible to biological researchers. The advantages of PGSFusion include the following:

1. Automatic identification of summary statistics. To simplify the process of uploading data for beginners, PGSFusion automatically recognizes the relevant data columns for analysis. To achieve this, we exhaustively cataloged many possible variable names and considered the expected characteristics of each variable type. The user can also manually specify the necessary parameters to address any incorrect assignments.
2. Comprehensive methods and reference panel. PGSFusion integrates 16 models belonging to four categories, including single-trait, multiple-trait, annotation-based, and cross-ancestry methods. When using multiple PGS models, a major challenge for beginners is that different methods require different formats for their reference panel. In response to this issue, PGSFusion offers ready-to-use reference panels from three ancestries for each method.
3. User-friendly interface. Each PGS method comes with its own unique hyper-parameters, which can be confusing for beginners to navigate. Based on our extensive experience, PGSFusion has simplified the parameter settings for each method to the maximum extent possible. For example, PGSFusion does not require any parameters for LDpred2.
4. Detailed epidemiological analysis and results. PGSFusion provides comprehensive analyses for epidemiological applications and generates high-resolution figures that are available for users to download. In addition, PGSFusion outputs the result of effect size in different PGS groups.
5. Extendible design. PGSFusion allows computational biologists to integrate novel PGS methods in the future to make them readily available to other researchers.

## Conclusions

PGSFusion is a new webserver that provides an easy-to-use, effective, and extensible solution for the construction and application of PGS. PGSFusion facilitates the selection of an optimal method for a given trait and requires no prior experience in running or developing PGS software to use. To further improve usability, PGSFusion can automatically infer the necessary inputs for PGS construction from summary statistics files. We demonstrate the benefits of PGSFusion through three comprehensive case studies covering a range of different application scenarios, data formats, and reference panels. By streamlining PGS construction and removing barriers of entry for biologists and epidemiologists, PGSFusion will further advance the research and clinical significance of PGS towards the objective of precision medicine.

## Methods

### Implementation

The frontend of PGSfusion is developed with ReactJs and styled with Antd, so users can effortlessly input gene fusion task details. An exciting addition to this feature is the automated population of separators and identifiers (e.g., rsID, *P*-value, Beta) upon uploading GWAS summary statistics files. This intelligent automation streamlines the user experience, enhancing efficiency and reducing errors. Meanwhile, on the backend, powered by Java and Spring Boot, a robust architecture ensures smooth task submission and execution. Leveraging in-memory caching and asynchronous task scheduling, our system optimizes resource utilization and responsiveness. As tasks are asynchronously dispatched, a dedicated timer periodically scans for task completion, marking them as finished once processed.

### Single-trait PGS methods

The algorithm of single-trait PGS methods is to select the independent SNPs with large effect size or to estimate the joint effect size of the whole genome. Following [13, 15], based on different distribution assumptions, PGSFusion classifies the 12 methods into six classes: i) no-model based: clumping and threshold (CT) [24, 29]; ii) one normal distribution: DBSLMM-lmm [23], LDpred2 [30], LDpred2-auto [31], PRSCS and PRSCS-atuo [32]; iii) two normal distributions with different variances: DBSLMM and DBSLMM-auto [23]; iv) point-normal mixture distribution: MegaPRS-BayesR [26]; v) latent Dirichlet process: SDPR [33, 34]; vi) Laplace distribution: lassosum2 [35]. Specifically, PGSFusion constructs the PGS for a specific trait and performs further analysis of it.

#### CT

CT relies on informed clumping and *P* value thresholding to select independent SNPs with large effect sizes. We provide three hyper-parameters for users to specify, including base window size, *R*^2^, and *P* thresholding. We recommended that the user used the default setting of the three parameters: i) window size: 50, 100, 200, and 500 kb; ii) correlation among SNPs: 0.01, 0.05, 0.1, 0.2, 0.5, 0.8, and 0.9; iii) number of *P* value thresholding on the log10 scale: 50. With the validation set consisting of 50 thousand individuals, CT selects the best parameter combination using the Pearson correlation coefficient (*R*^2^) (for quantitative traits) or area under the curve (AUC) for binary traits. PGSFusion uses the *bigsnpr* R package (v.1.12.2) to fit the CT model.

#### DBSLMM, DBSLMM-auto, and DBSLMM-lmm

DBSLMM relies on a mixture of two normal distributions to assume the effect size of each SNP. To select independent SNPs with large effect size by PLINK (v.1.90b6.9), DBSLMM set the window size to 1000 Kb, the correlation between SNPs to 0.2, and the *P*-value thresholding to 1×10^-6^ [25]. With the SNP heritability estimated by linkage disequilibrium score regression (LDSC), DBSLMM should select the best heritability from 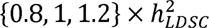 by the validation set [13, 23]. DBSLMM-auto automatically sets the SNP heritability and *P*-value threshold choice of 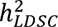 and 1×10^-6^, respectively. In addition, based on the LMM assumption, DBSLMM-lmm only uses the reference panel without parameter tuning. PGSFusion uses the *DBSLMM* (v.0.3) software to fit the three models.

#### lassosum2

lassosum2 relies on the penalty regression to estimate the SNP effect size with two hyper-parameters, including the shrinkage parameter *δ* and penalty parameter *λ*. Following [36, 37], based on the validation set, lassosum2 tunes four choices of *δ* to be 0.001, 0.01, 0.1, and 1, and 30 choices of *λ* evenly spaced on a log10 scale between *λ_min_* and *λ_min_*/100 with *λ_min_* in each chromosome. PGSFusion uses the *bigsnpr* R package (v.1.12.2) to fit the lassosum2 model.

#### LDpred2 and LDpred2-auto

LDpred2, a new version of LDpred, relies on updating the joint effect size of each SNP and proportion of causal SNPs with the LD structure by Gibbs sampling [30, 38]. With the SNP heritability estimated by linkage disequilibrium score regression (LDSC), LDpred2 uses the validation data to tune 63 (21×3) parameter combinations, including a sequence of 21 values from minimum value to 1 on a log-scale for the proportion of causal SNPs, and three choices of 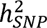 as 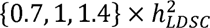 [30]. Adding a parameter modeling negative selection, LDpred2-auto uses 50 different polygenicity parameters to fit the LDpred model and regards the mean of beta as the estimator [31]. Note that we set the burn-in number and iteration number as 500 and 500, respectively. PGSFusion uses the *bigsnpr* R package (v.1.12.2) to fit the two LDpred2 models.

#### MegaPRS-BayesR

MegaPRS-BayesR assumes a mixture of a point mass at zero and three normal distributions on the SNP effect sizes and heritability varying based on the MAF of the SNP [26, 39]. Performing a pseudo cross-validation with 10% samples for testing, MegaPRS-BayesR inherently specifies 35 candidate hyperparameter combinations of *π*_1_, *π*_2_, *π*_3_, and *π*_4_ . PGSFusion uses the *LDAK* software (v.5.2) to fit the MegaPRS-BayesR model and the LDAK-Thin model [26, 40].

#### PRSCS and PRSCS-auto

PRSCS relies on continuous shrinkage (CS) prior, including global scaling parameter, *ϕ*, and local, SNP-specific parameter, *ϕ*, to allow for the shrinkage consistent with the genetic architecture [32]. Based on the validation set, PRSCS sets the two hyper-parameters, *a* and *b*, for the three-parameter beta prior on the shrinkage factor, τ = 1/(1+ *ϕϕ*), to the default values to 1 and 0.5, respectively, and selects *ϕ* from {10^-6^, 10^-4^, 0.01,1} with the best prediction performance. PRSCS-atuo, the automatic version of PRSCS, estimates ¢ from the reference panel alone. To reduce the computational time, PRSCS set the burn-in number and iteration number as 1000 and 500, respectively. PGSFusion uses *PRScs* software to fit the two PRSCS models.

#### SDPR

SDPR relies on the latent Dirichlet process prior on the SNP effect size [33]. Similar to DPR, the SDPR model uses Markov Chain Monte Carlo (MCMC) algorithm to estimate posterior inference using summary statistics and its corresponding reference panel [33, 34]. PGSFusion uses the *SDPR* software to fit the SDPR model.

### Multiple-trait PGS Method: *mtPGS*

Constructing PGS for a target trait of interest leverages multiple traits relevant to the target trait. mtPGS, an extension of DBSLMM, relies on the joint model to estimate the effect size of two traits with a high genetic correlation [41]. Following [23], mtPGS specifies the SNP correlation and the *P*-value threshold as 0.2 and 10^-6^ in default, respectively. Based on the genetic correlation provided by the *GECKO* R package (v.1.0), PGSFusion uses the *mtPGS* software (v.1.0.0) to estimate effect sizes for the target trait under a bivariate modeling framework using the reference panel.

### Annotation-based PGS method: *AnnoPred*

Assuming that SNPs located in functional areas are highly enriched for signals, AnnoPred sets functional annotations to assign empirical priors to SNPs [42]. AnnoPred provides four “tier” choices, including the smoothed GenoCanyon annotation, seven tissue-specific GenoSkyline annotations, and 66 cell-type-specific GenoSkylinePlus annotations [38, 42]. Using the validation set, AnnoPred considers 11 choices for the proportion of causal variants, including 1, 3×10^-1^, 1×10^-1^, 3×10^-2^, 1×10^-2^, 3×10^-3^, 1×10^-3^, 3×10^-4^, 1×10^-4^, 3×10^-5^, and 1×10^-5^. PGSFusion uses the *AnnoPred* software to fit the model.

### Cross-ancestry PGS methods

Leveraging summary statistics of EUR GWAS in large-scale, cross-ancestry PGS improves prediction performance in other ancestries, such as AFR and EAS. PGSFusion includes three new and best-performance methods: PRSCSx [43], SDPRX [44], and XPASS [45].

#### PRSCSx

Assuming CS coupling across ancestries, PRSCSx, an extension of PRSCS, jointly estimates the SNP effect in each ancestry with two summary statistics and two reference panels [32, 43]. Different from PRSCS, PGSFusion provides PRSCSx without tuning ¢ and set the burn-in number and iteration number to 100 and 500, respectively. PGSFusion uses the *PRScsx* software (v.1.1.0) to fit the model.

#### SDPRX

SDPRX, an extension of SDPR, specifies a joint distribution as the prior on the effect sizes in two ancestries to be both null, population-specific, or shared across two ancestries and assumed the proportions of the variance component matrix following the Dirichlet distribution. With trans-ethnic genetic correlation estimates by PopCorn (v.0.0.1), PGSFusion uses the *SDPRX* software to fit the model.

#### XPASS

Assuming that the genetic correlation largely remains in different ancestries, XPASS sets the effect size as a bivariate normal distribution and obtained the analytic estimation [45]. Apart from the reference panel for both populations, PGSFusion also treats the top 20 principal components (PCs) in EUR and the top 5 PCs for EAS/AFR as covariates. PGSFusion uses the *XPASS* R package (v.0.1.0) to fit the model.

### Epidemiological applications in UKBB cohort

PGSFusion conducts comprehensive and in-depth analyses of PGS prediction performance and its interactions, with additional analyses for summary statistics without UKBB individuals (Fig. 1). The calculation and its visualization in PGSFusion not only provide a detailed assessment of PGS performance but also shed light on more nuanced differences in genetic risk and its interaction with key epidemiological factors.

For performance evaluation, PGSFusion provides the following functions: i) overall performance to investigate the predictiveness of PGS for the corresponding trait; ii) genetic subgroup performance to show variation in predictiveness between different genetic risk groups; and iii) covariate subgroup performance to show variation in predictiveness between groups stratified by a particular covariate. Generally, summary statistics are regressed against the effects of sex, age, and the top PCs, so that the performance evaluation reports the total effect from both the PGS and these covariates. For quantitative traits, we use the model in Equation (1):

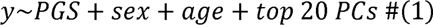

where *y* is the analyzed trait. PGSFusion then plots the joint density between the PGS and the trait, and evaluates predictiveness using *R*², Cor(*y*, *ŷ*)^2^ along with its 95% confidence interval (CI). For binary traits, we use the model in Equation (2):

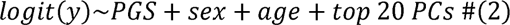

where *y* is the analyzed trait. PGSFusion uses a receiver operating characteristic (ROC) curve to measure overall predictiveness in terms of the area under curve (AUC) as *AUC*(*y*, *ŷ*) along with its associated 95% CI.

To compare the predictiveness of a PGS at different levels of genetic risk, PGSFusion categorizes individuals by their genetic risk quantile into equally sized groups and displays a forest plot where each point and error bar indicates the risk effect and its 95% CI for a group. Similarly, for subgroups stratified by a covariate, PGSFusion uses a forest plot to show prediction performance for each group (e.g., female and male for sex), along with the *P*-values from a Wilcoxon test for statistical significance.

For joint analysis, PGSFusion provides two kinds of analysis. To identify interactions overall, PGSFusion uses a bar plot to display the genetic risk effect (effect size divided by odds ratio) for the subgroups defined by a specific covariate, along with the 95% CI and *P*-value for the significance of the interaction with that covariate. PGSFusion continues to adjust for sex, age, and top PCs when modeling interactions. For example, the interaction between PGS and sex is given by:

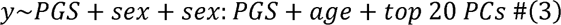

where *sex: PGS* indicates the interaction term. To identify interactions within subgroups of individuals categorized by a covariate or effect quantile, PGSFusion utilizes a forest plot to display the effect size for different covariates in each subgroup.

### Linkage disequilibrium (LD) matrix of reference panel

PGSFusion construct the LD matrix of the reference panel for three ancestries using the 1000 Genomes Project (1000GP) [27], with different strategies and different data formats to be compatible with each PGS method. We categorize the reference panels for the methods into three classes, with details for each method in Table 1. First, CT, AnnoPred, LDpred2, and MegaPRS-BayesR model the LD structure as block matrices of up to a maximum distance in base pairs (e.g., 2 Mb for AnnoPred) or genetic distance (e.g., 3 cM for LDpred2 and MegaPRS-BayesR). Second, following [47], PRSCS, DBSLMM, SDPR, lassosum2, mtPGS, PRSCSx, and XPASS model LD using a block-diagonal matrix. Third, SDPRX accounts for admixture when modeling LD. For the methods that fall under the first two strategies, PGSFusion provides separate LD matrices for each of the three ancestries in 1000GP, whereas for SDPRX, we provide LD matrices for EUR-EAS and EUR-AFR admixtures.

### Data processing for validation and test sets

After processing UKBB data with the same procedures as in our previous publications [13, 15, 23], we obtained 337,129 EUR, 6,442 AFR, and 3,538 EAS individuals. For the methods requiring hyperparameter selection, we randomly selected validation set I. For the two epidemiological applications of prediction performance and joint analysis, we randomly selected two sets of EUR individuals as validation set II and test set. Each set contained 50,000 individuals, of which 25,000 are female [46]. Specifically, the three sets had no overlapping individuals. Validation set I was used to select hyper-parameters for the PGS methods CT, LDrped2, DBSLMM, PRScs, lassosum2, and AnnoPred. These six methods in PGSFusion were only suitable for summary statistics estimated from EUR populations and excluding UKBB individuals. For all 16 methods, validation set II was used to estimate the effect size assigned to PGS and other covariates. In addition to evaluating in-ancestry prediction on the test set containing 50,000 EUR individuals, we also evaluated cross-ancestry prediction using all of the AFR and EAS individuals as additional test sets.

To ensure reliable results, we conducted stringent quality control (QC) on the quantitative and binary traits we model. For quantitative traits, we selected traits that were *h*^2^ > 0.1 in the EUR population. Binary traits included both disease (i.e., AD) and non-disease traits (i.e., type I baldness). Among diseases, we included self-reported traits (fields 20001 and 20002), diagnoses indicated by ICD10 codes (fields 41202 and 41204), and diseases that are specifically defined (fields 2453 for cancer, 2443 for diabetes, and 6150 for vascular/heart problems). For diseases and other binary traits, we kept traits where i) the ratio of cases to control is greater than 1:500, and ii) 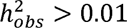 . After QC, we selected 125 quantitative traits, including standing height, BMI, and platelet count, and 41 binary traits (Additional file 2: Table S6 and S7). Following [15] and to ensure the scalability of all PGS methods, we only used SNPs contained in phase 3 of HapMap (HM3).

To ensure proper evaluations using the summary statistics, we fitted a linear regression for quantitative traits or logistic regression for binary traits in validation set II to estimate the effects of the constructed PGS, the top 20 genotype PCs, and covariates including sex and age, then applied the estimated effect size to the test set. Specifically, for quantitative traits, we performed a *z*-score transformation on the phenotypes of the two validation sets and test set. Note that PGSFusion also provided the additional ability to handle summary statistics estimated by sex-specific or age-specific analysis.

### Summary statistics of three case studies

We demonstrated the utility of PGSFusion through three case studies involving GWAS summary statistics collected from different ancestries and different traits. In Case Study 1, we showed PGS construction using all single-trait methods, followed by epidemiological analyses using the UKBB cohort. For this study, we used AD summary statistics from PGC (https://vu.data.surfsara.nl/index.php/s/l7aiRr1UEgdoJfZ), which were estimated from 71,880 cases and 1,036,225 controls [7]. In Case Study 2, we investigated the cross-ancestry performance of PGS constructed using three cross-ancestry methods and compare their performance to three corresponding single-ancestry methods applied to EAS individuals from the UKBB cohort. This case study used standing height summary statistics from EAS (participants from BBJ: https://humandbs.biosciencedbc.jp/files/hum0197/hum0197.v3.BBJ.Hei.v1.zip) and EUR (participants from GIANT Consortium: https://portals.broadinstitute.org/collaboration/giant/images/0/01/GIANT_HEIGHT_Wood_et_al_2014_publicrelease_HapMapCeuFreq.txt.gz) ancestries with 165,056 individuals in BBJ [48] and 253,288 individuals in GIANT [49], respectively. In Case Study 3, we evaluated the performance of a multiple-trait method and compare it with a single-trait methods. This case study used summary statistics for two correlated traits with EUR ancestry, which were weight from GIANT (http://www.broadinstitute.org/collaboration/giant/index.php/GIANT_consortium_data_files) estimated from 60,586 EUR males [50] and high-density lipoprotein (HDL) cholesterol (https://csg.sph.umich.edu/willer/public/glgc-lipids2021/results/sex_and_ancestry_specific_summary_stats/HDL_INV_EUR_HRC_1KGP3_others_MALE.meta.singlevar.results.gz) estimated from 67,9743 EUR males [51]. In every case study, we evaluated the performance and computational cost of the selected PGS methods and conducted further analyses using the features of PGSFusion.

## Ethics approval and consent to participate

Not applicable.

## Availability of data and materials

Code for each method and UKBB processing is on Github (https://github.com/biostat0903/PGSFusion/tree/main). All relevant data is available through the PGSFusion website (http://www.pgsfusion.net/).

## Authors’ contributions

C.C., S.Y., and P.H. conceived the project and designed the pipeline. S.Y., X.Y., X.J., Z. L., and T.M. implemented and evaluated the software. S.Y. and C.C. wrote the manuscript.

## Acknowledgement

This study has been conducted using UK Biobank resource under Application Number 1144904. We are grateful to the participants and study staff of UK Biobank.

