## Supplementary Figures for "*PGSFusion* streamlines polygenic score construction and epidemiological applications in biobank-scale cohorts"

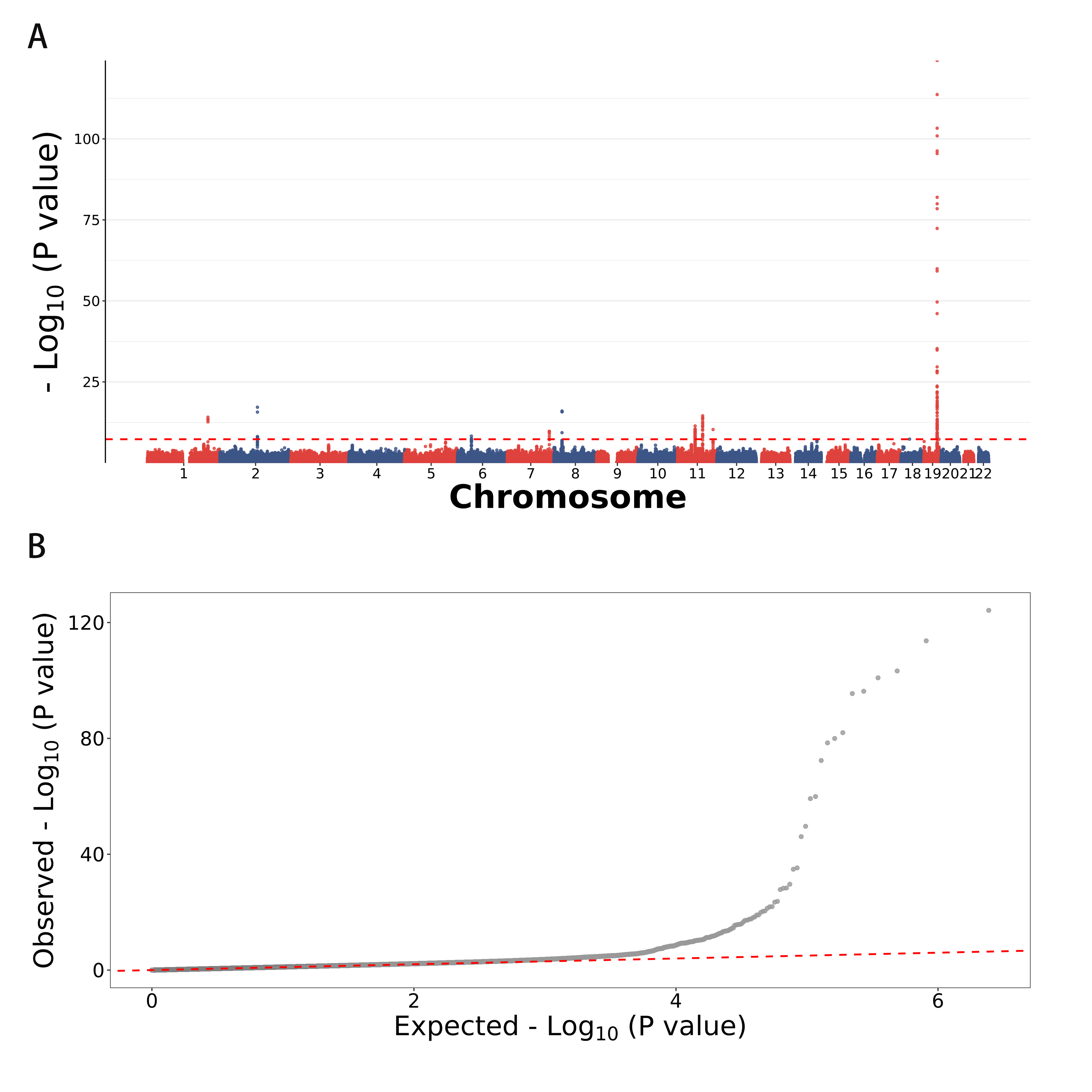

**Fig. S1 Summary for AD summary statistics.** A) Manhattan plot. B) qqplot.

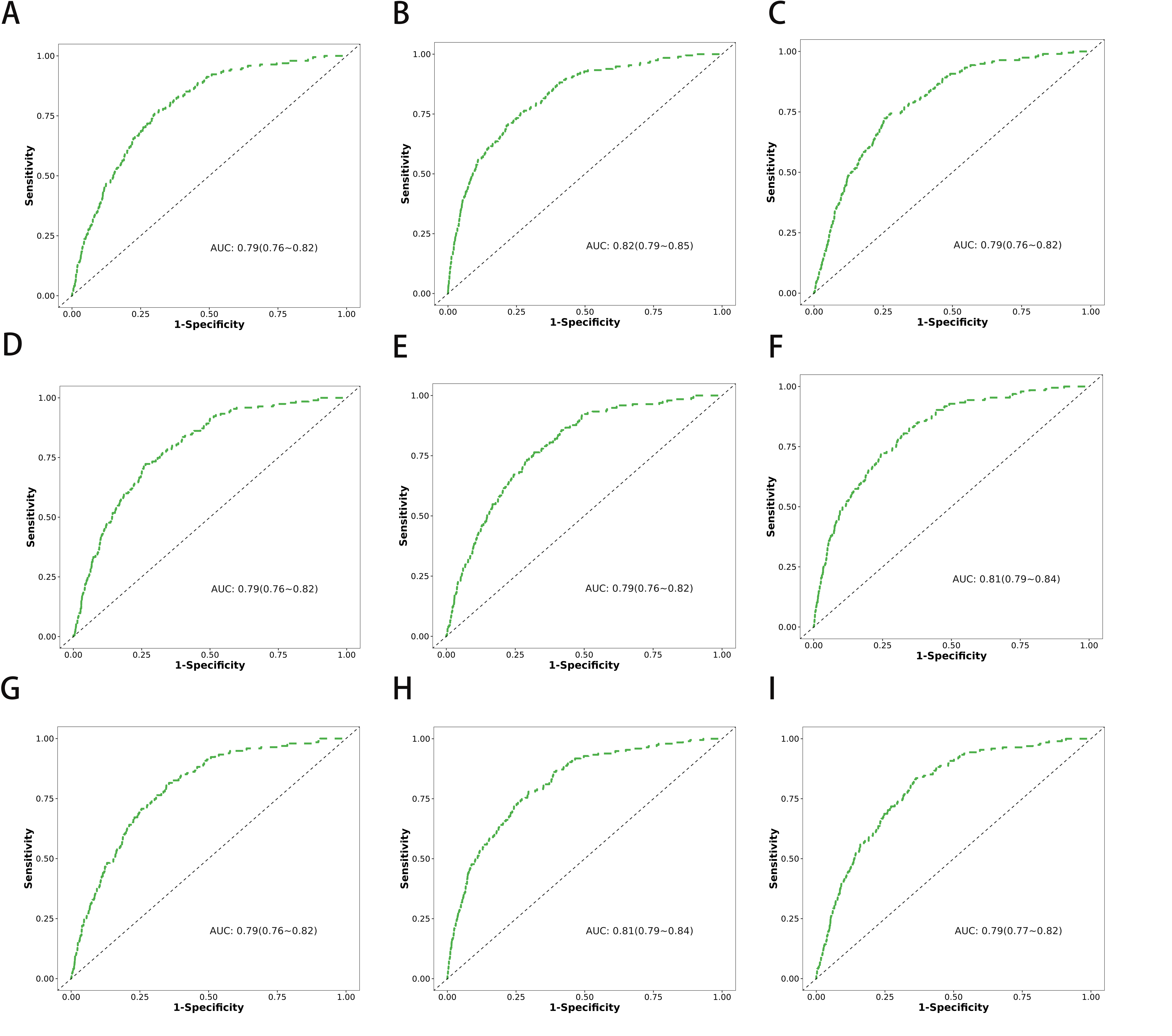

**Fig. S2 ROCs of the nine remaining methods.** The nine methods are CT, lassosum2, LDpred2, LDpred2-auto, MegaPRS, PRSCS, PRSCS-auto, and SDPR.

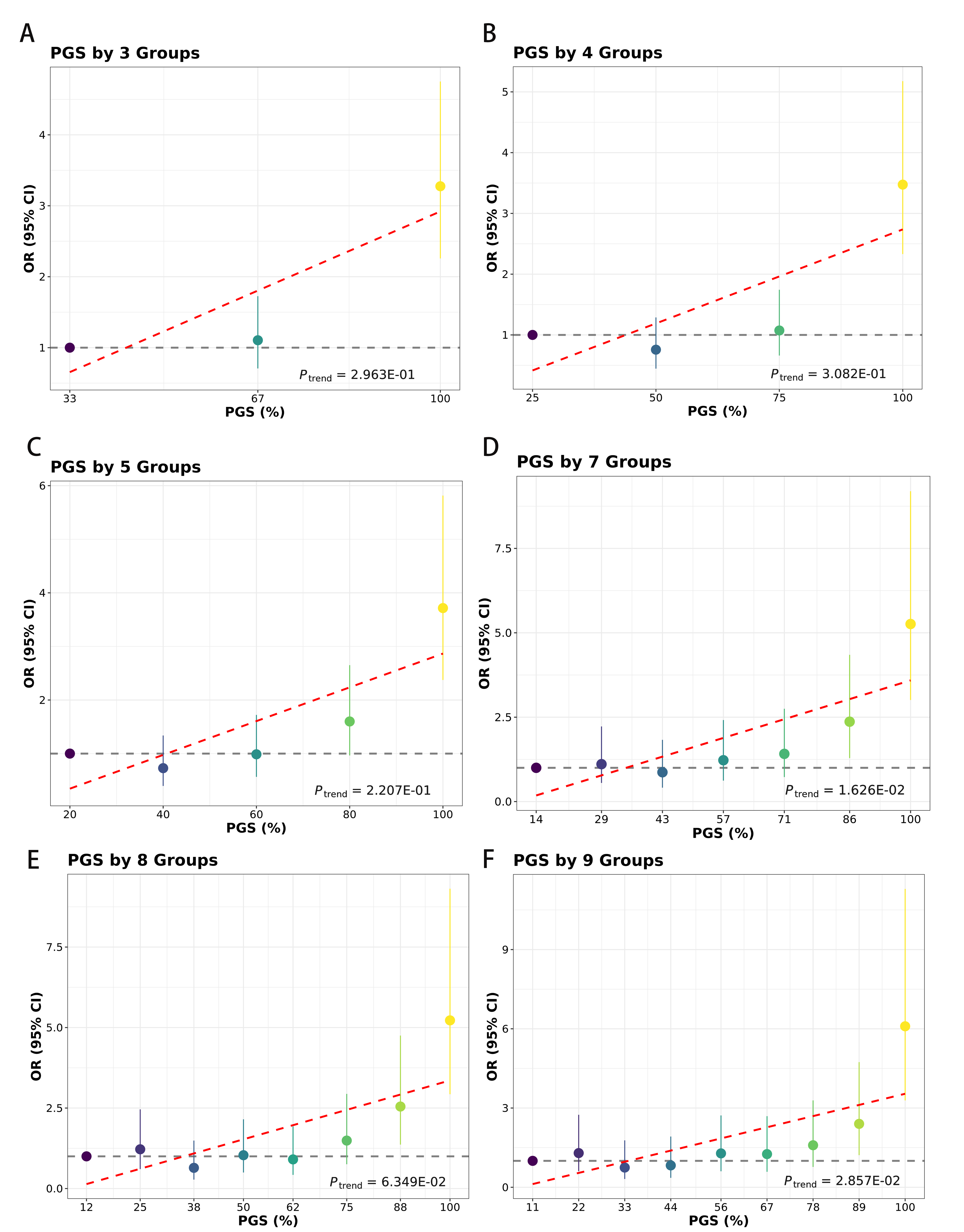

**Fig. S3 Effect size of different PGS groups with six settings of group number.** Individuals in the EUR test sets were divided into equal groups according to PRS, and the ORs of each group were compared with those in the lowest group. Error bars are 95% CIs.

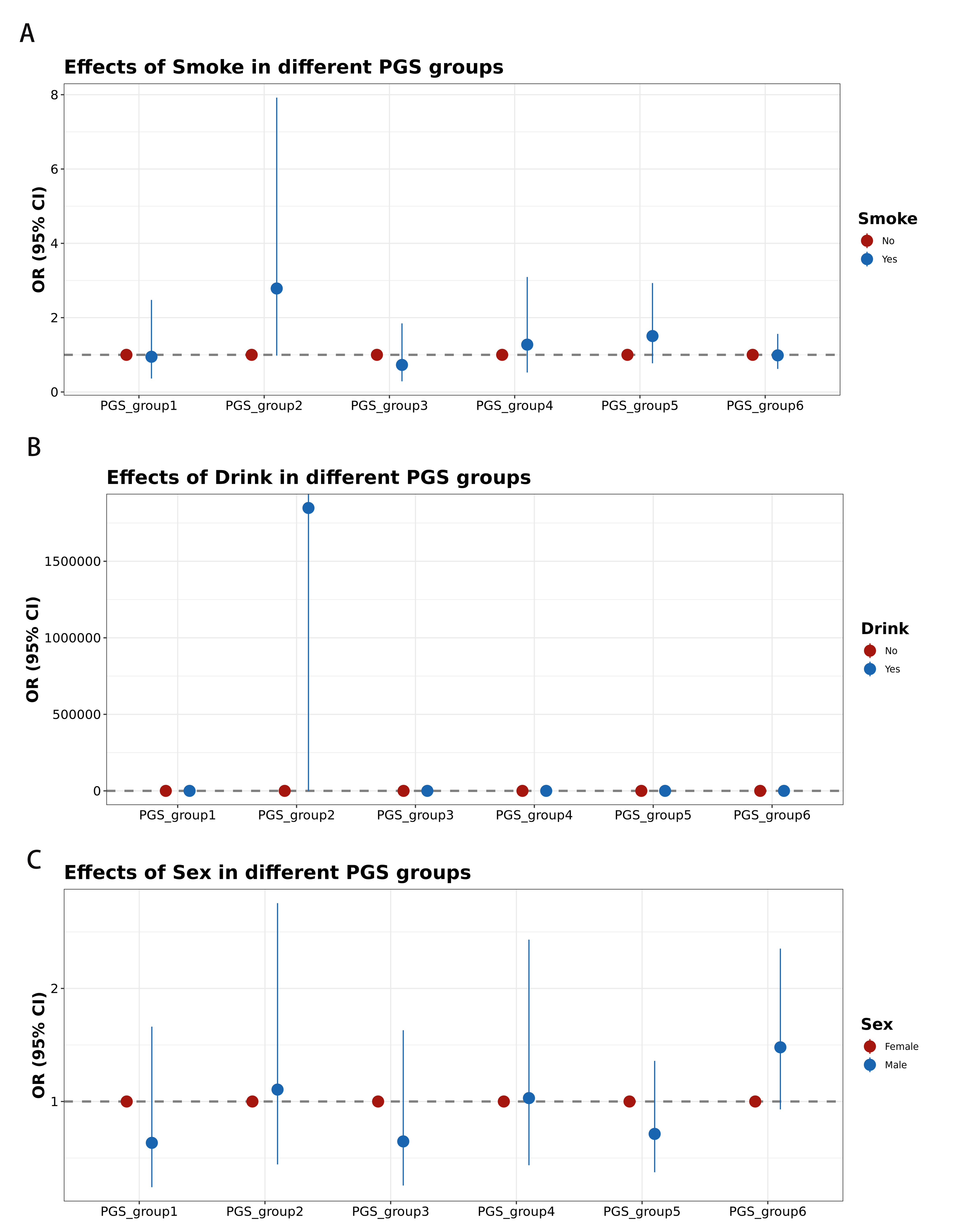

**Fig. S4 The effect size of smoking, drinking and sex in different PGS groups.** A) Effect size of smoking in six PGS groups. B) Effect size of drinking alcohol in six PGS groups. C) Effect size of sex in six PGS groups.

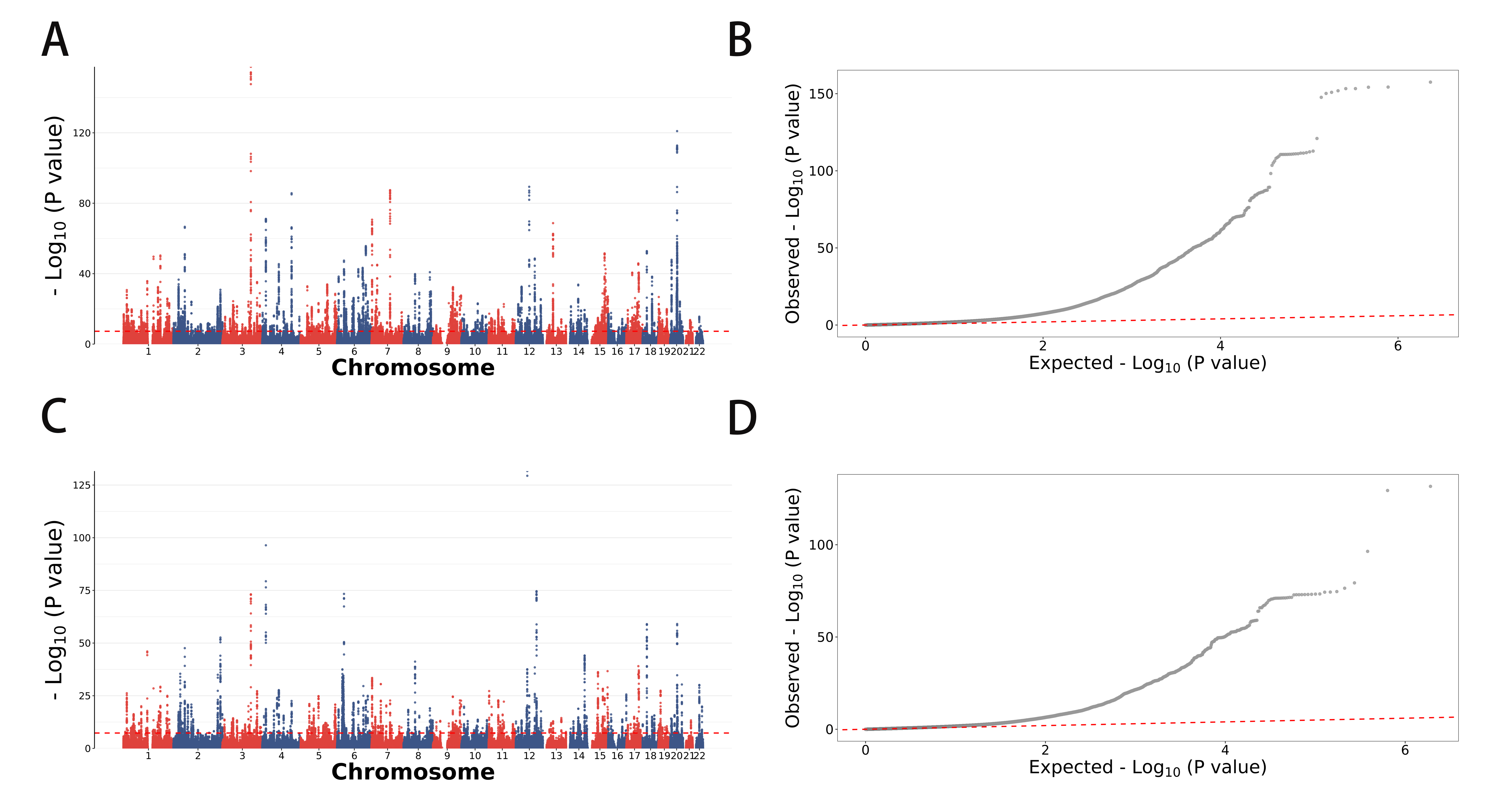

**Fig. S5 Summary for height summary statistics in GIANT and BBJ.** A) Manhattan plot for GIANT. B) qqplot for GIANT. C) Manhattan plot for BBJ. B) qqplot for BBJ.

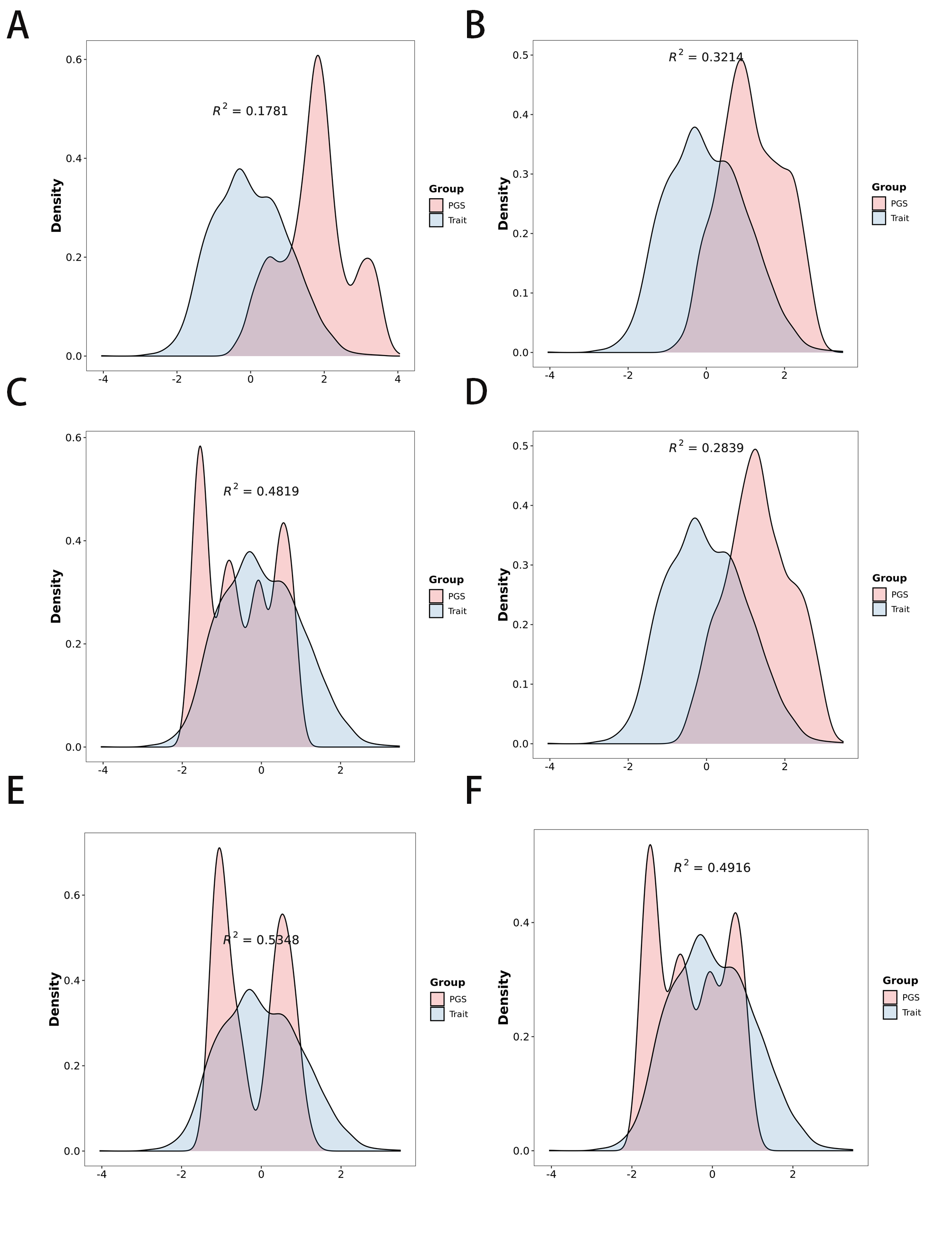

**Fig. S6 ROCs of the six methods for Case Study 2.** The six methods are DBSLMM-auto, PRSCS-auto, PRSCSx, SDPR, SDPRX, and XPASS.

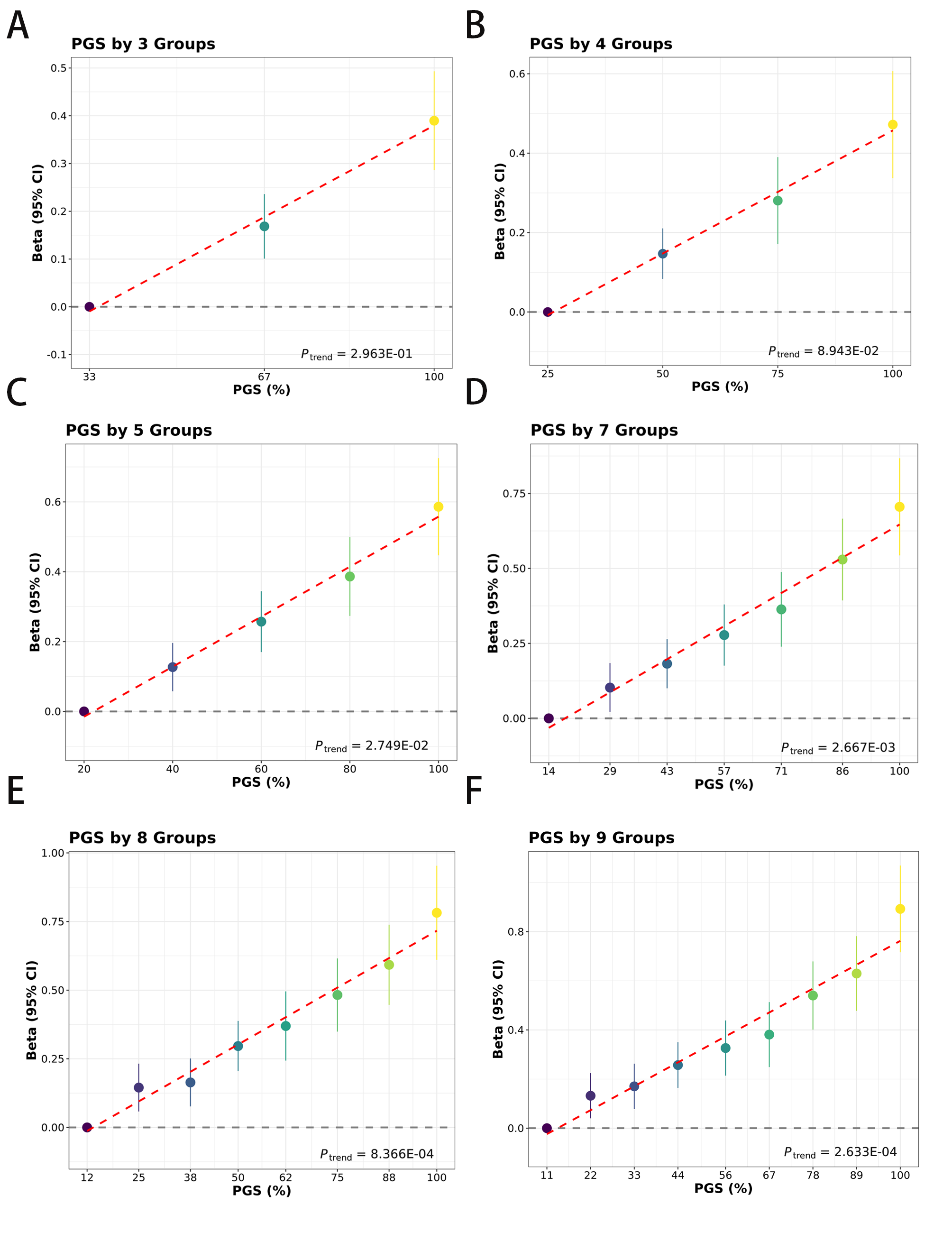

**Fig. S7 Effect size of different PGS groups with six settings of group number.** Individuals in the EAS test sets were divided into equal groups according to PRS, and the Beta of each group were compared with those in the lowest group. Error bars are 95% CIs.

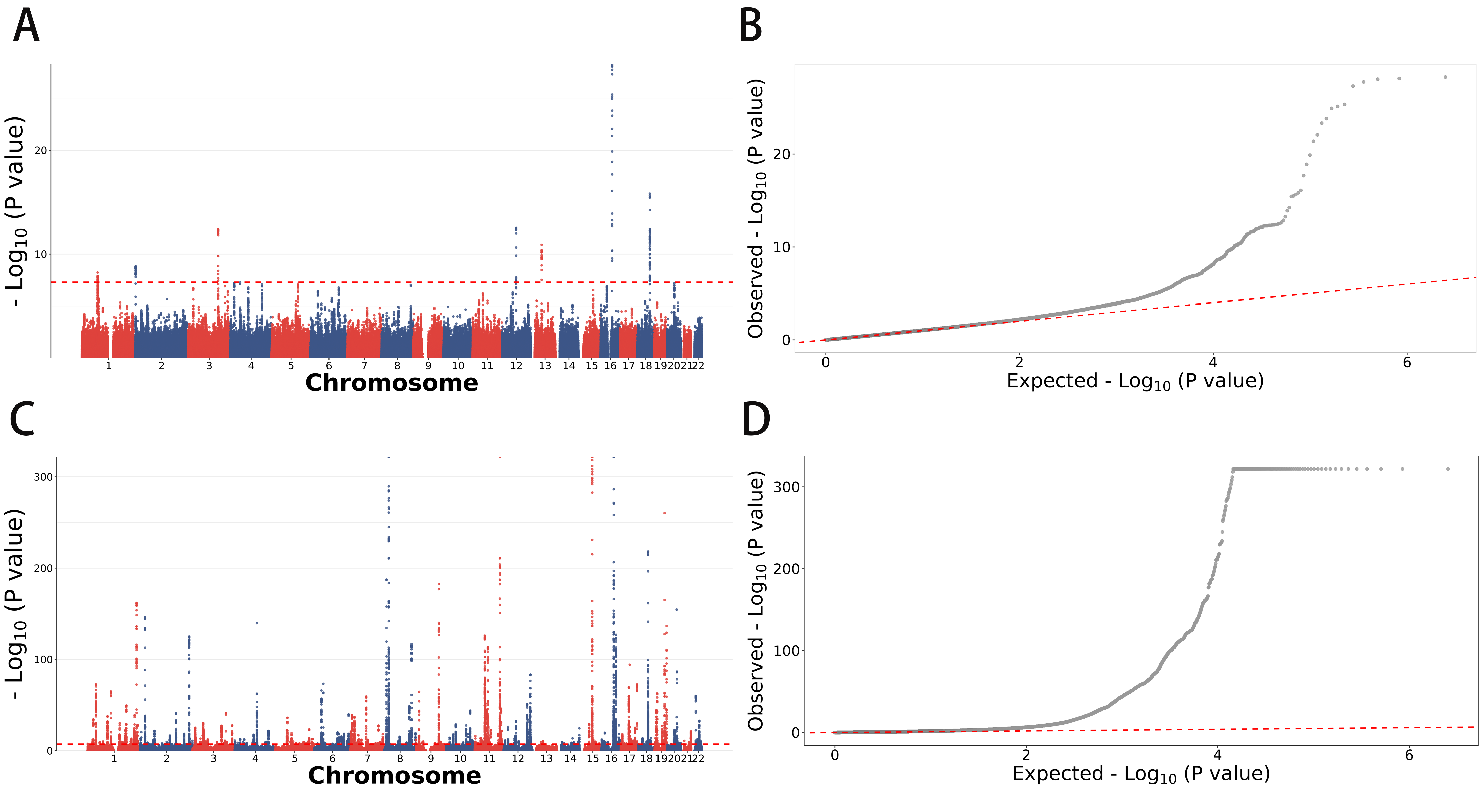

**Fig. S8 Summary for weight and HDL summary statistics.** A) Manhattan plot for weight. B) qqplot for weight. C) Manhattan plot for HDL. B) qqplot for HDL.

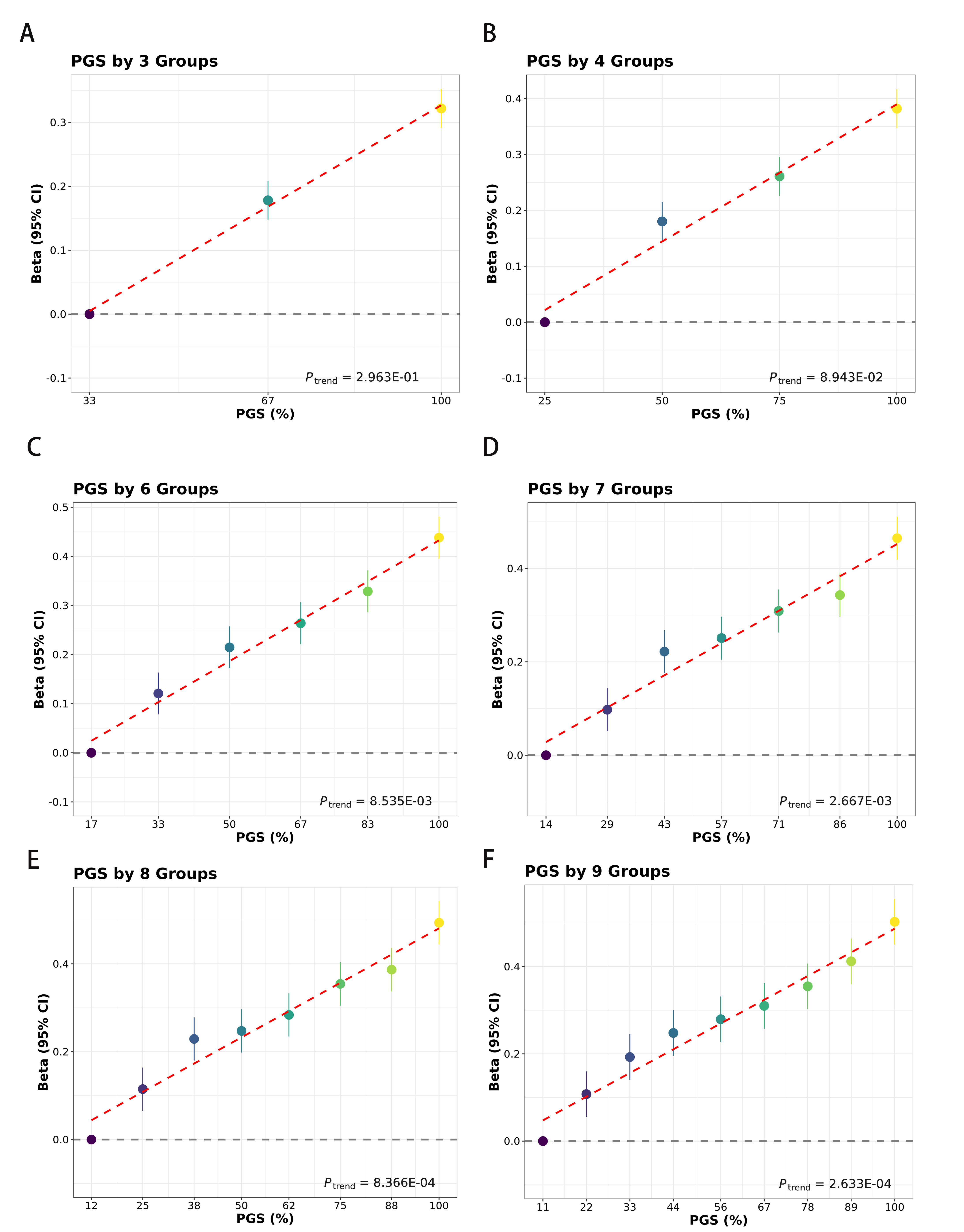

**Fig. S9 Forest plots for mtPGS in different group number settings for mtPGS.** Individuals in the EUR test sets were divided into equal groups according to PRS, and the Beta of each group were compared with those in the lowest group. Error bars are 95% CIs.

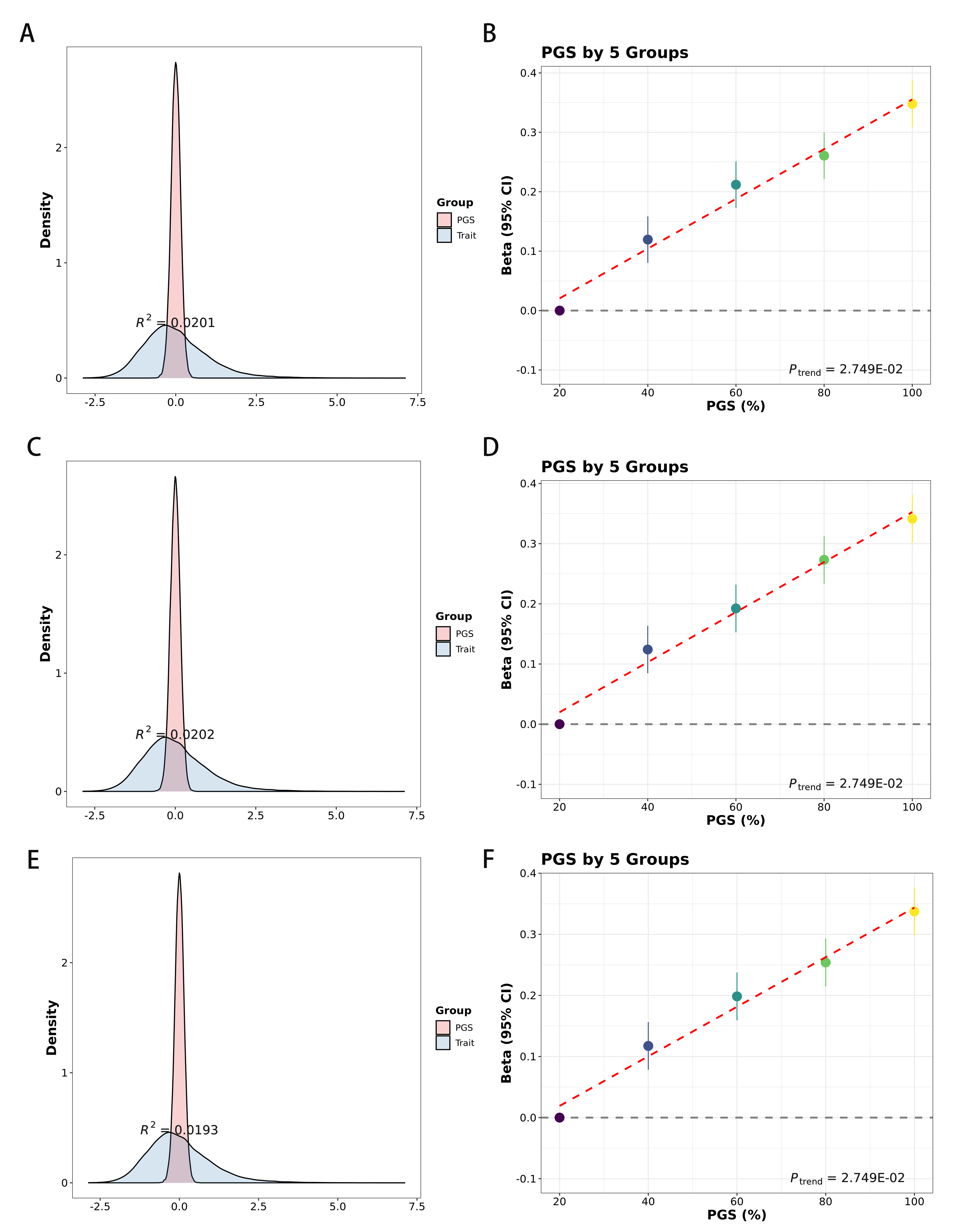

**Fig. S10 Summary of overall prediction performance for DBSLMM, DBSLMM-LMM and DBSLMM-auto.** A) Density plot for the test set and weight PGS for DBSLMM. B) Individuals in the test sets were divided into five PGS groups for DBSLMM, and the ORs of each group were compared with those in the lowest 1st group. C) Density plot for the test set and weight PGS for DBSLMM-LMM. D) Individuals in the test sets were divided into five PGS groups for DBSLMM-LMM, and the ORs of each group were compared with those in the lowest 1st group. E) Density plot for the test set and weight PGS for DBSLMM-auto. F) Individuals in the test sets were divided into five PGS groups for DBSLMM-auto, and the ORs of each group were compared with those in the lowest 1st group.

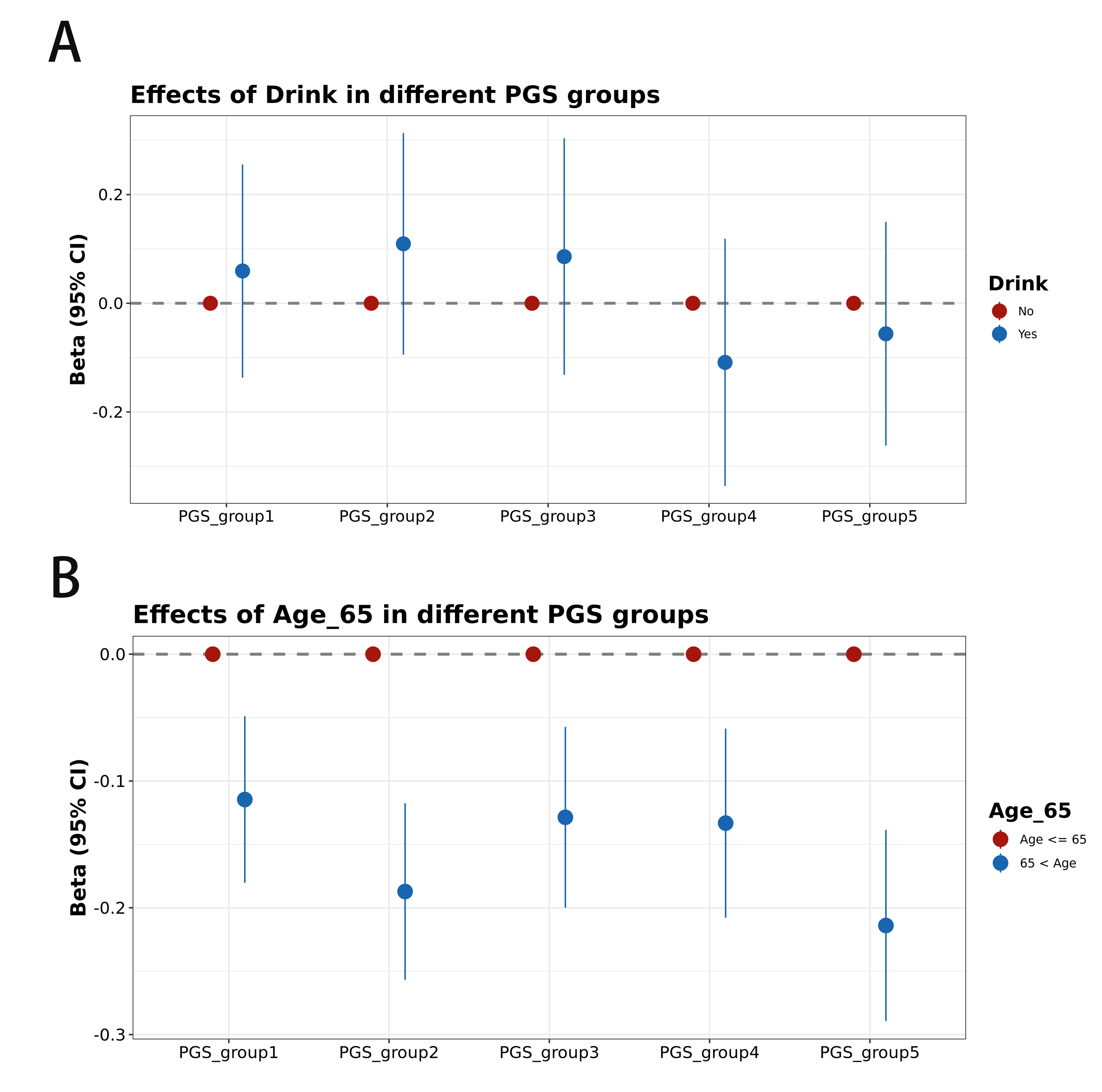

**Fig. S11 Summary for the effect size of smoke, drink and age in five PGS groups.** A) Forest plot displays the effect size of alcohol (No vs Yes) in the five PGS groups. B) Forest plot displays the effect size of age (<65 vs >65) in the five PGS groups.
